# Machine-learning strategy for high error tolerance in image-based digital molecular assays

**DOI:** 10.1101/2025.08.14.670347

**Authors:** Darren B. McAffee, Qiang Hu, Assame Arnob, Hung-Jen Wu, Jay T. Groves

## Abstract

There is a significant global health need to translate more *in vitro* diagnostic tests (IVDs) from clinical laboratories to field-based applications, including point-of-care (POC) and self-administered test formats. These applications typically require smaller sample sizes, limit the extent of sample processing and measurement capabilities, and introduce greater handling variability. Error tolerance is one of the most critical factors for successful field-based assay design. Here, we examine machine-learning (ML) strategies to enhance the error tolerance of image-based nanoparticle immunoassays. Random dispersions of nanoparticles were imaged in microliter sample volumes, and images were processed to determine analyte concentrations based on nanoparticle appearance. Assay performance was characterized using two common blood diagnostics: C-reactive protein (CRP) and S.CoV-2 IgG. We compare the results from a conventional image analysis, a hybrid ML-conventional approach based on pixel segmentation, and a full end-to-end image regression using a targeted regularization strategy. Training images for the full image regression approach required only a single label for training – the analyte concentration – eliminating the need for labor-intensive pixel-level labeling. Ultimately, the fully ML-based analysis significantly improved dynamic range, sensitivity, and reproducibility in high-error settings, including direct measurements performed in whole blood.

## Introduction

The digitization of immunoassays—shifting from bulk signal measurements to individual particle-by-particle analyses—is a rapidly advancing trend in biomarker detection^1^. This analog-to-digital transition in molecular detection mirrors, in many ways, the analog-to-digital transition in the electronics industry. Similarly, it holds great promise to achieve higher sensitivity, precision, and especially error resistance in molecular diagnostics. Techniques like Single Molecule Array (Simoa)^2^ or Single Molecule Counting (SMC)^3^ assays achieve ultrahigh sensitivity by isolating individual target molecules within femtoliter-sized wells or droplets, respectively, thereby digitizing the assay. Existing digital assay methods still require precision-engineered well arrays, droplet generators^4^, or high-end imaging setups^5,6^ to minimize background noise from non-specific binding, optical scatter, and image artifacts—all of which can severely impact accuracy. Error tolerance isn’t an automatic benefit of molecular assay digitization. However, high pixel count images of particle-based assays include vast amounts of information beyond the analyte detection signals themselves. This additional information can be used to identify and filter noise and error sources that would otherwise be inexorably blended in the assay readout^7^.

While sensitivity improvements have historically driven innovation in immunoassay development, reliability and error tolerance are often more critical for POC and self-testing^8–13^. Many clinically relevant protein biomarkers are present at picomolar (pM) or higher concentrations^14^, which are readily detectable by many immunoassay formats. Diagnostic errors arising from pre-analytical factors, such as sample contamination and improper handling, and post-analytical factors, such as ambiguous readout interpretation, present greater challenges to field-based assay development^15–18^. The real-world coefficient of variation (CV) of an assay is generally of greater importance than the limit of detection (LOD) established in controlled settings^8,19,20^. There is a substantial need for technology advancements that increase assay error-tolerance and simplicity. In this study, we explore ML-driven error filtering to increase error tolerance and assay performance in image-based digital immunoassays.

We examine biomarker assays based on the clustering behavior of gold nanoparticles (AuNPs)^21^, but the strategies employed are broadly applicable to other signal reporters such as shaped plasmonic nanoparticles^22,23^, colloidal particles^24–27^, fluorophores^6^, or many other reporter types^28^. Functionalized AuNP assays for CRP and S.CoV-2 IgG were performed on microliter sample volumes of serum and whole blood. Direct imaging of nanoparticle clustering by darkfield microscopy provides measurable improvements over bulk spectroscopic readouts, increasing both sensitivity and dynamic range. While standard image processing algorithms^29^ functioned adequately in limited circumstances, they were often brittle and required meticulous sample preparation to minimize error.

We sought to improve image-based digital assay performance using two different ML strategies. First, an ML-assisted strategy based on pixel segmentation to filter out image artifacts prior to conventional image analysis offered improvements, but still required tedious training. Finally, by coupling a modified ResNet image classification model^30^ with a targeted regularization strategy, we efficiently trained models using only a single label—analyte concentration—eliminating the need for pixel-level labeling. Effective training could be accomplished with ~10,000 images and a single GPU. This end-to-end image regression approach differs substantially from conventional or hybrid ML analyses in that it requires no user-defined criterion to interpret images. All aspects of the image containing useful information are utilized, providing substantial performances improvements in high error settings. With full ML analysis, clinical laboratory level performance (e.g. equivalent to ELISA) was achieved with an extremely simple mix-and-measure workflow on whole blood microsamples (**Figure 1a**). These findings highlight the potential of image-based digital immunoassays to leverage rapidly developing ML capabilities. The resulting reduction in hardware and workflow complexity while maintaining high diagnostic accuracy is a crucial step toward developing next-generation POC diagnostics that are robust, widely accessible, and ready to meet global health needs.

**Figure 1.**
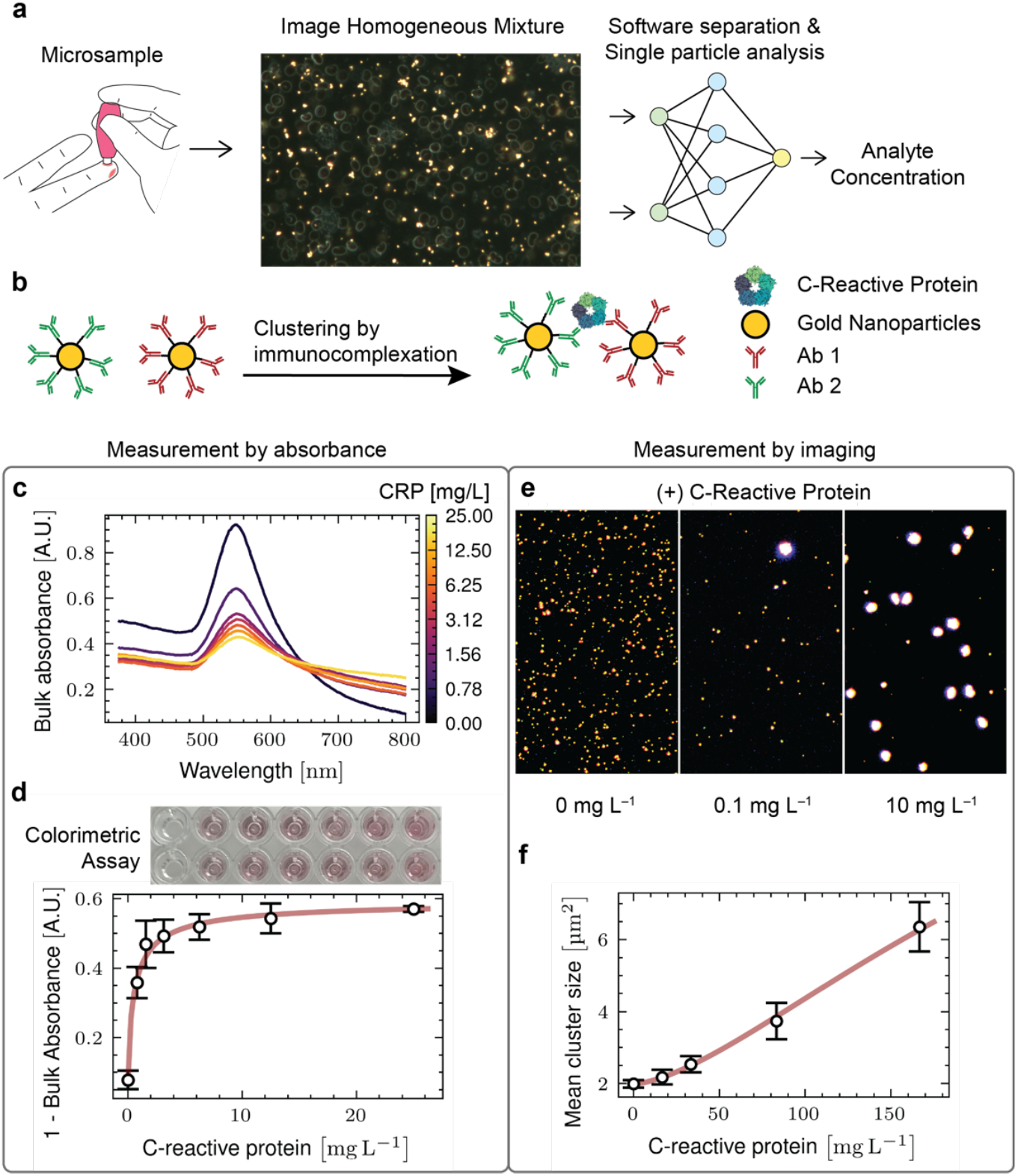
(a) Schematic of the simplified workflow of a digital assay using random particle arrays of gold nanoparticles imaged in-situ with sample, (b) Design of particle aggregation assay towards C-reactive protein, (c/d) Traditional absorbance based methods for inference of analyte concentration, (c) Spectra of samples with increasing CRP concentrations with associated decrease in 550 nm absorbance and simultaneous red-shifting, (d) Titration of CRP concentration against spectral shifts, (e) Example images using 20x darkfield to image the nanoparticle aggregates in the presence of increasing CRP concentration, (f) Titration of CRP concentration against quantification of aggregate size.

## Results

### Image-based readout with conventional analysis

Biomolecular assays based on functionalized AuNPs are widely used in diagnostics, including lateral flow assays (LFAs)^31^ and bulk solution colorimetric assays^21^. In these formats, capture agents linked to the AuNP (e.g. antibodies, aptamers, nanobodies, etc.) bind the target analyte. Two capture agents binding distinct epitopes on the target will lead to crosslinking or localization of the particles in an analyte-dependent manner (e.g. sandwich immunoassay). Alternatively, multivalent targets can crosslink particles with even a single capture agent^32^. In LFAs, signal is produced as enhanced optical density by concentrating nanoparticles at a designated test strip location whereas signal in solution-based assays is detected as color or light scattering changes caused by nanoparticle oligomerization. LFAs and solution-based assays are generally analyzed by bulk optical readout, in which individual particles are not resolved—only an average optical response from all particles is read. However, individual nanoparticles can be readily imaged by simple techniques such as dark-field microscopy^33–35^. Image-based assays employ microscopy to directly resolve the entire dispersion of nanoparticles, offering the possibility to individually evaluate each nanoparticle or cluster. Image analysis steps are then employed to interpret nanoparticle features from the images (e.g. mean nanoparticle cluster size, color, or brightness), which can be related to analyte concentration^36,37^. An important advantage of image-based assay readout is the possibility to filter out various forms of error (e.g. dust, bubbles, pre-aggregated particles etc.) from the final analyte concentration determination based on definable image characteristics^7^.

To assess the performance advantages of image-based readout, we first examine a classic two-antibody sandwich AuNP immunoassay for CRP. AuNPs (80 nm diameter) were conjugated with two CRP-specific antibodies (**Figure 1b**). Clustering of anti-C-reactive protein (α-CRP) particles by the CRP analyte results in a red-shifted absorption-scattering peak, which can be seen by eye and read quantitatively by UV-Visible spectroscopy^21^. Assays were run by incubating α-CRP particles with CRP in plasma or whole blood at varying concentrations. As CRP concentration increased, the 555 nm peak in the absorption-scattering spectrum decreased (**Figure 1c**). Quantification of the 555 nm absorbance peak magnitude provides a robust readout for the assay but it has a limited dynamic range, due partly to larger aggregates falling out of solution as well as the fact that spectral changes with increasing aggregate size become less pronounced (**Figure 1d**). When AuNPs from the same sample solutions are directly imaged (**Figure 1e)**, numerous aggregates are visible and quantifiable with straightforward image processing techniques ^29^. Mean cluster size provides a robust linear response to CRP concentration that extends more than 30 times beyond the saturation limit of the colorimetric assay (**Figure 1f**). The imaging readout readily covers the clinical concentration range for CRP diagnostics (<1mg/L to 200 mg/L) using the same particles that failed to cover this range in bulk spectroscopic readout.

### ML-assisted analysis based on pixel segmentation

While conventional image analysis based on predefined criterion can be used to perform a rough separation of image artifacts from functional nanoparticles, a tremendous amount of information about the sample remains unused with such methods. Here we examine an ML-assisted image analysis strategy based on pixel-wise segmentation to computationally separate genuine signal from artifact. Various ML algorithms can learn to do pixel-wise segmentation and there exist open-source software packages enabling facile training^38,39^. In this hybrid format, ML is first used to separate out error sources in images, then a conventional readout (e.g. mean cluster size) is performed on the ML-filtered images to infer analyte concentration.

We characterize this ML-assisted analysis on AuNP assays for S.CoV-2 IgG levels in patient samples. AuNPs were functionalized with a single capture protein, S.CoV-2-RBD, here referred to as α-IgG(SCoV-2-RBD) particles (**Figure 2a**). These α-IgG(SCoV-2-RBD) particles exhibited robust aggregation in the presence of immunoglobulins (G and E type) specific to S.CoV-2-RBD (**Supplemental Figure 2a and 2b**). Training images were generated using the pixel classification workflow in *ilastik* ^38^ and used to train random forest (RF) ML algorithms (see Methods). Although the resulting RF models could be trained quickly with minimal data, they lacked robustness and failed to generalize beyond 8–10 images. To address this limitation, we built an ensemble of seven RF models, each trained on 5–6 annotated images (≈ 50,000 particles per image) representing a range of sample types (serum, blood, and PBS) as well as varying degrees of defocus. This ensemble approach helped maintain model coherence and captured greater variability. The combined output — 45 fully pixel-labelled high-resolution (4K) images — served as the training dataset for a U-net ^40^. We applied standard data augmentation techniques from the PyTorch library to enhance segmentation performance (see Methods for detailed descriptions). As a result, the U-net significantly outperformed the directly generated RF models in terms of generalization and was able to rapidly segment images into relevant classes such as background, nanoparticles, dust, and debris. After pixel-wise classification on the blood samples and removal of erroneous objects **(Figure 2b**), the mean cluster size of the nanoparticles can be quantified and mapped to analyte concentration (**Figure 2c**).

**Figure 2.**
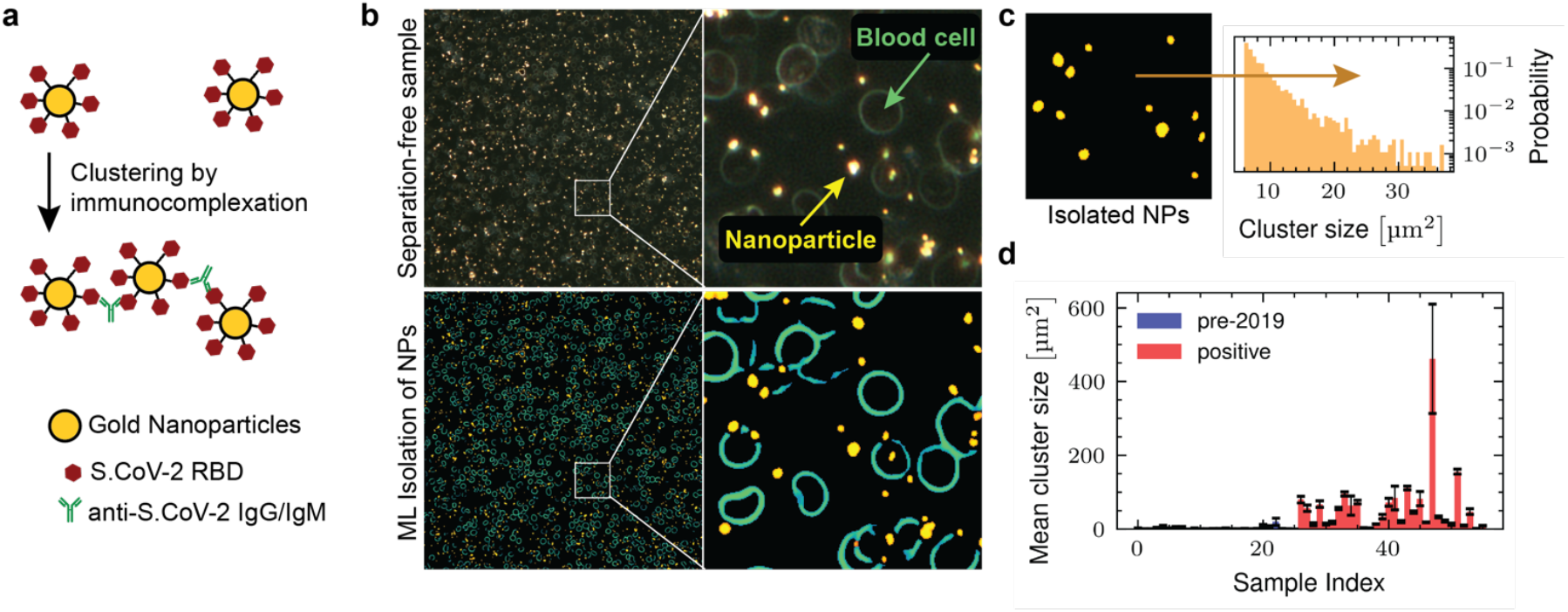
(a) Design of gold nanoparticle assay - aggregation of particles induced by the presence of antibodies against the RBD of S.CoV-2. (b) Example images using 20x darkfield to capture nanoparticle aggregation in whole blood samples. The probability maps for different classes (gold nanoparticle, cell, background) produced by the segmentation model are displayed below while the original image is on top. (c) Post-processing of cluster size shows an approximate power law distribution, (d) Quantitation of cluster size on biobank sample from two patient groups: one 3 weeks convalescent from S.CoV-2 and the other group are true negative (pre-2019) samples.

We used this trained pixel segmentation algorithm to quantify IgG(S.CoV-2-RBD) in patient samples. A total of 55 samples were tested, including 25 true negative (pre-2019) and 30 true positive (PCR positive, 3-week convalescent) samples. The patient samples were imaged in a relatively high-throughput open air microwell format, which also introduces high levels of error including bubbles, dust, as well as optical distortions due to the meniscus of the well (**Supplemental Figure 2c**). Despite these challenges, the digital molecular assay robustly classified positive COVID-19 samples (**Figure 2d**). The positive prediction value was 77% = 23/30 and is comparable to the seroconversion rates (79% and 88%) detected with gold standard clinical laboratory methods^41,42^.

This use of pixel-segmentation for error filtering followed by traditional analysis still relies on predetermined particle features (e.g. cluster size, color, brightness, etc.) in the final image analysis step to infer analyte concentration. A vast amount of analyte-dependent information contained in the images is still left unutilized by such an approach. We hypothesized that an end-to-end image regression approach, which maps a full sample image to analyte concentration, could potentially learn which features of the nanoparticles are most important while also learning to ignore optical distortions and other artifacts.

### End-to-end image regression for full ML assay interpretation

We developed a full end-to-end image regression approach for assay interpretation by training a modified ResNet-18 model^30^ on 10,000 images labelled only with analyte concentration (**Figure 3a**). Approximately 55 non-overlapping images were recorded on each of 182 samples consisting of various concentrations of S.CoV-2 IgG doped in human serum. The labels corresponded to a titrated dataset with 7 concentrations (0, 16, 64, 256, 1024, 4096, and 16384 ng/uL S.CoV-2 IgG). The dataset was split into testing and training groups (164 training samples, 18 test samples). Separation by sample (instead of just image) proved important to ensure that model generalization comes from nanoparticle properties rather than sample specific artifacts; several types of sample error span multiple images (e.g. nanoparticle adherence patterns or extended scratches on the substrate).

**Figure 3.**
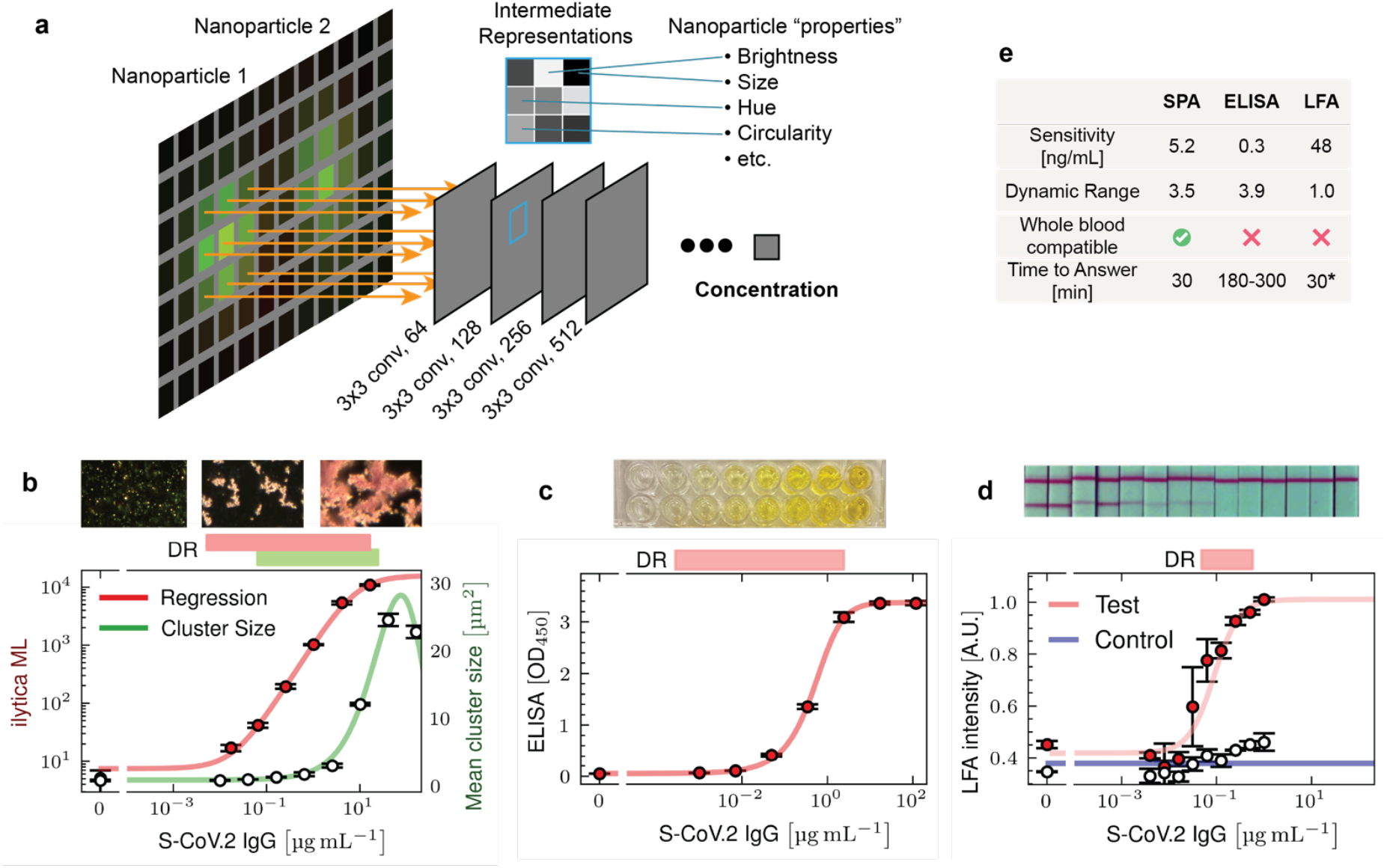
(a) Schematic of how the end-to-end machine learning model works. The image is processed by several convolutional layers. Deeper layers can capture more abstract properties of the particles and allow surrounding context to influence outcomes, (b) Titration of S-Cov.2 IgG against the output of the end-to-end ML mod el compared to the output of the hybrid segmentation + post-processing model, (c) Titration of S-Cov.2 IgG using a commercial ELISA kit. (d) Titration of S-Cov.2 IgG using a Universal Lateral Flow Kit with the same particles used in the imaging assay (b). (e) Table comparing the single-particle analysis (SPA) method from (b), the ELISA method from (c) and the LFA method from (d).

For our initial attempt, the final layer of the ResNet model consisted of a collection of neurons, corresponding to the number of classes the model is intended to predict (e.g. the 7 trained concentrations). Using this classification approach, we trained the model to predict which labelled concentration the unknown sample image belonged to. While effective, this required additional interpolation methods to infer sample concentrations between the classifications. We next tested a direct regression architecture, where the final layer of the ResNet-18 had a single output neuron intended to only predict the log of the analyte concentration. This single-neuron output strategy had significantly better performance. Concentration estimates were made from the mean prediction of 20 images from the sample.

The results from application of the regression model to the testing samples, with known concentrations of S.CoV-2 IgG spanning the titration range, are shown in **Figure 3b (red trace)**. Results from the hybrid ML approach (pixel-segmentation followed by traditional image processing) are also plotted in **Figure 3b (green trace)** for comparison. While the hybrid approach does remove sources of error, it is clearly limited by only analyzing a handful of properties of the nanoparticles. While there is no direct map between human and machine interpretation, we speculate that the full regression model relies on a spectrum of image properties that shifts with analyte concentration. For example, properties such as particle density and color ratios are likely more important at lower analyte concentrations, while other properties such as aggregate size and void area are more heavily weighted at higher concentrations. The model is not restricted to following one parameter across the entire detection range. We quantified the assay sensitivity using a fitted Hill function in conjunction with the measurement error around the blank and lowest concentration to estimate the limit of detection^43^. The regression model achieved near-ELISA capabilities on a first attempt with a sensitivity of 34.7 pM (5.2 ng/mL) and a dynamic range of 3.5 orders of magnitude—beating the hybrid ML model by more than an order of magnitude in both characteristics.

For benchmark comparisons, similar S.CoV-2 IgG titration results were obtained from traditional ELISA (clinical lab-standard) and LFA (POC-standard) assays. The ELISA was performed from a commercial testing kit (Thermofisher) and achieved a typical sensitivity of 2 pM (0.3 ng/mL) with a dynamic range of 3.9 orders of magnitude (**Figure 3c**). For a direct comparison to LFA, we created an in-house LFA using the same α-IgG(SCoV-2-RBD) particles as were used in the image-based experiments using a universal LFA kit (Abcam). LFA test strips were imaged and the relative intensity of test and control strips were assessed using a laboratory scanner (see Materials and Methods). While the LFA achieved a respectable sensitivity of 0.32 nM (48 ng/mL), it had a poor dynamic range with barely 1 order of magnitude (**Figure 3d**), limiting its use as a quantitative assay. Moreover, our use of a laboratory scanner to quantify the LFA results likely improved its performance beyond typical POC applications.

In addition to core assay metrics like sensitivity and dynamic range, there are other important distinctions between the overall formats of the assays. For blood-based testing, both ELISA and LFA usually require plasma/serum separation steps to avoid signal degradation from whole blood. In contrast, an image-based ML approach can be used directly on whole blood, reducing the hardware complexity required. LFA is noted for its rapid turnaround, and image-based digital assay workflows have shown similar turnaround times. Both LFA and image-based digital assays are limited by the primary incubation time between capture and target proteins, which is governed by reagent on and off rates with target. ELISA typically has a significantly more complex workflow, taking 3-5 times longer. These comparisons are summarized in **Figure 3e**.

### Sensitivity of cross-linking digital assays

Detection limits in image-based readout of nanoparticle assays are intrinsically at the single molecule level^34^, offering the potential to vastly surpass bulk readout sensitivity. Practically, the overall assay sensitivity limit in an imaging readout is determined by the number of definitive analyte molecular detection events and the ability to distinguish these from nonspecific signals. Analysis of more particles effectively increases the number of trials, thereby reducing noise through counting statistics. In contrast, for bulk measurements such as ELISA, increasing the sample volume does not inherently improve sensitivity. In fact, it can dilute the analyte concentration or introduce additional background noise, potentially lowering the signal-to-noise ratio and making it more difficult to detect low-abundance targets. In the following, we provide a general analysis relating measurable physical properties of the nanoparticle sensors to corresponding assay sensitivity over a broad range of digital assay implementation conditions. The results detailed below illustrate how assay sensitivity can be adjusted, with almost no limit, by adjusting the number of particles utilized in the assay. These analyses apply generally to digital molecular assays. ML error filtering strategies increase both the number of useable particles as well as the fidelity of signal discrimination.

To assess sensitivity scaling laws for digital assays we consider two critical assay features: particle polydispersity and intrinsic optical signal strength. While this analysis focuses on particle clustering assays, it is generally applicable to any droplet or bead digital assay format. We approximate the starting distribution (**Figure 4a**) to consist of monomers with a Gaussian size distribution, and pre-existing aggregates (dimers, trimers, etc.) are modeled by convolving this distribution with itself. The quality of the starting particle dispersion is thus characterized by the monomer-dimer separability (σ) and the fraction of pre-existing aggregates (λ), see **Figure 4b and 4c**. The key question to determine analyte detection limits then becomes: when analytes crosslink particles from some baseline distribution, how many particles must be sampled to reliably detect a difference? Using a Monte Carlo method coupled with likelihood ratio testing we determined how this detection limit depends on σ and λ. For the experimental assays described in this study, 5 μ*L* serum samples were mixed with 20 μ*L* particle solutions for a final testing mixture of 25 μ*L* containing ≈ 300 million AuNps and analyte at 1/5 original sample concentration. The inexpensive AuNps used here have σ/μ = 0.38 and λ = 0.15 (**Figure 4a**). The assay achieved 34.7 pM by sampling ≈ 1 million particles. Higher precision nanoparticle manufacturing and purification processes can produce particles with significantly lower polydispersity^44^, which can dramatically improve assay sensitivity. However, an even larger sensitivity improvement is achieved by lowering the number of particles in the sample incubation (increasing the fraction of particles that capture analyte). Reducing particle number comes at a cost of incubation time and potentially the use of re-concentration methods.

**Figure 4.**
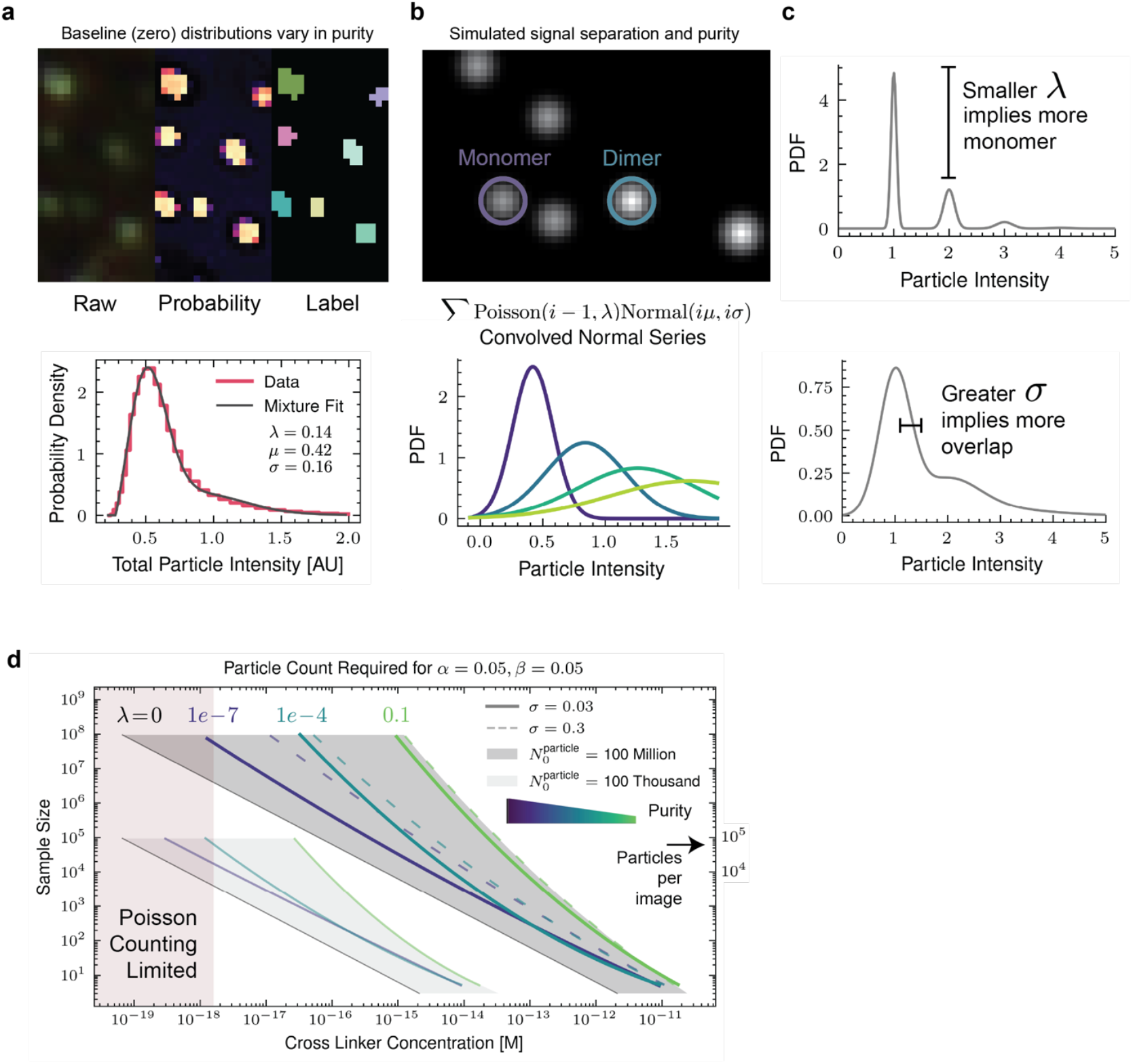
(a) Example image of gold nanoparticles with zero analyte. A histogram of particle intensities is shown below along with a model fit. The model used in the histogram is described by (b) A Poisson weighted mixture model of a series of convolved normals is used to approximate a starting particle distribution. Simulated images showing the intensities of monomer and dimer is shown on top. (c) Graphs demonstrating how the different parameters of the mixture model influence total intensity distributions of the aggregates, (d) Line plot of scaling laws (sample size required for a given limit of detection) determined by a monte carlo approach.

We calculated scaling curves that map the number of particles that must be sampled to achieve desired analytical detection limits for various particle properties (σ, λ) (**Figure 4d**). The assay particles used here approximately correspond to the light green line. Assay sensitivity can be improved using higher precision manufacturing and purification processes to reduce polydispersity, which can get down to λ = 1 × 10^−4^, albeit at higher cost. The leftmost black lines represent fundamental performance limits with perfect particles and discrimination distribution (σ = 0, λ = 0) – this corresponds to the stochastic sensing limit that no assay can surpass. The overlap between monomer and dimer intensity values (σ) was not as impactful as the monodispersity of the initial particle distribution (low λ). For example, even if monomer and dimer intensities have 25% overlap in their distributions, subfemtomolar concentrations can be reliably detected with 100 images worth of particles. We note that atto-or zeptomolar detection limits can be achieved by reducing the total number of particles (**Figure 4d**). This increases the fraction of particles that capture analyte, but comes at a cost of longer incubation times.

## Discussion and Conclusion

Here we examined two different ML strategies for improving image-based assay performance: a hybrid approach combining pixel segmentation with traditional image processing and an end-to-end deep learning model that directly mapped images to analyte concentrations. While both methods improved over conventional assays, the latter significantly outperformed the hybrid approach in terms of sensitivity and robustness. The hybrid approach effectively removed background noise and extracted useful nanoparticle properties, but it was inherently limited by predefined feature selection. The end-to-end classification approach, by contrast, leveraged all available image features and dynamically learned which properties were most indicative of analyte concentration. This ability to self-correct and learn from diverse sources of noise led to notable improvements in dynamic range, sensitivity, reproducibility, and general tolerance to error and unskilled handling. ML training was accomplished with moderate data sets that can be easily collected in a manner of hours with simple laboratory equipment and a single GPU. It is entirely practical for specific ML algorithms to be trained for every particle batch on a wide range of imaging platforms or in a made-to-order format.

This study introduces an ML-driven approach to digital molecular assays that significantly enhances overall performance while reducing hardware complexity. By leveraging image-based analysis and deep learning, we demonstrate a system that outperforms both ELISA and LFA in key metrics such as sensitivity, dynamic range, and robustness to error. This advancement represents a critical step toward more accessible, reliable, and scalable diagnostic solutions, particularly for POC and self-administered testing applications. As the field of digital molecular diagnostics continues to evolve, the integration of computational intelligence with molecular assays holds immense potential to transform healthcare by making high-quality diagnostics more widely available.

## Methods

### Gold Nanoparticle Conjugation and Characterization

Carboxylated gold nanoparticles (AuNPs) of 80 nm diameter (NanoComposix SKU: AUXR80-5M) were functionalized with antibodies or proteins for specific target recognition. For C-reactive protein (CRP) assays (α-CRP particles), two distinct monoclonal antibodies against CRP – anti-C2 (abcam ab17452) and anti-C6 (abcam ab244707) were conjugated to AuNPs via amine-NHS ester conjugation. For the α-IgG(SCoV-2-RBD) particles detecting SARS-CoV-2 immunoglobulin (IgG/IgE), recombinant S.CoV-2 receptor-binding domain (RBD) protein (Abcam AB273065-1003) was conjugated to AuNPs to form a single-capture format. Nanoparticle conjugation was confirmed by analyte-specific aggregation under darkfield microscopy. Functionalized particles were stored in 0.1x PBS at 4 C until use.

### Colorimetric Bulk Assays

CRP detection using bulk absorbance was performed by incubating α-CRP AuNPs with recombinant CRP (Abcam AB283925-1003) at various concentrations (0.78 – 166 mg / L). After incubation for 1 hour at room temperature, UV-Visible absorbance spectra were collected using a spectrometer. Absorbance at 555 nm was quantified and normalized to control samples lacking CRP. Peak shifts and aggregation signatures were assessed by monitoring changes in the spectra across 500–700 nm.

### Imaging and Sample Preparation

Samples were imaged using an Olympus BX43F equipped with an SC180 CMOS camera under darkfield illumination conditions. For static imaging, 5 uL of sample was mixed with 2 µL of nanoparticles in 18 uL 0.1x PBS buffer (25 uL total) and incubated for 30 minutes. Then, 6 uL of incubated mixture was placed onto a standard 75×25mm microscope slide and covered with a 1 cm^2^ coverslip. In assays involving human blood, 1 µg of recombinant IgG-α-S.CoV-2 RBD (Thermofisher MA535939) was spiked into 1 mL of whole blood (Fisher Sci NC1570874). Some imaging was done in open-air microwell formats were used to intentionally introduce imaging artifacts such as dust, bubbles, and meniscus distortion for testing ML robustness.

### Hybrid Pixel-wise Segmentation and Traditional Image Processing

The pixel segmentation model we used was a Unet trained on a collection of images, each labelled by one of 7 random forest classifier (RFC) models created using ilastik. Training images for the RFCs included nanoparticle clusters and background elements such as blood cells, bubbles, and other artifacts. The RFC models used by ilastik perform well on homogeneous datasets but fail to generalize across broader sets of images. As such, sets of 5-10 images were grouped to maintain RFC performance and 7 separate RFC models were trained spanning the diversity of images in the dataset. We note that training RFCs with several classes (background, particles, blood cells, bright debris, stratch, etc.) outperformed a simple background/foreground approach. A Unet was subsequently trained on the pixel classified images using standard data augmentation techniques. Once the Unet was trained, images were segmented by their predicted probability for pertaining to the particle class. Post-processing of the probability maps included morphological filtering, size exclusion, watershed filtering, and shape-based filtering to separate distinct objects. Cluster metrics such as mean area, brightness, and shape descriptors were extracted using the scikit-image library for Python.

### End-to-End Deep Learning Model Training

An end-to-end image regression model was developed using a ResNet-18 backbone via the PyTorch library: `torchvision.models.resnet18.` A total of 10,000 darkfield images from 182 samples were used to train the model, with sample concentrations titrated across 7 levels (0 to 16384 ng/µL). Data was split into training (164 samples) and testing (18 samples) sets by sample, ensuring no data leakage due to repeated, albeit non-overlapping, imaging of the same specimen. The model was trained on unnormalized images. The final regression layer output a single scalar corresponding to the natural logarithm of concentration, the zero concentration was labelled as −0.6931 while the second lowest concentration (16 ug/mL) was labelled as 2.773. Model performance was validated on a separate test set using the mean prediction from 20 images per sample.

### Clinical Samples and Assay Validation

Clinical human serum samples were obtained from Precision for Medicine (PFM) and included 25 pre-2019 samples (true negative controls) and 30 PCR-confirmed convalescent COVID-19 samples (true positives). PFM collects samples with a FDA 21 CFR part 11 compliant biospecimen inventory system. Two images of each specimen were analyzed in triplicate wells (6 total images) per sample and imaged after direct addition of nanoparticle mixture as described in above in sample preparation. Performance metrics such as positive predictive value and false positive rate were calculated from the model outputs.

### ELISA and Lateral Flow Assays

The ELISA comparison was done with a Human SARS-CoV-2 Spike (trimer) IgG ELISA Kit from Thermofisher (BMS2325). Assays were performed following manufacturer instructions. Absorbance was read at 450 nm using a microplate reader. The limit of detection and dynamic range were calculated from the standard curve. A lateral flow assay was assembled in-house using a Universal LFA kit (Abcam AB270537-1001) and the above described α-IgG(SCoV-2-RBD) AuNPs as reporters. Test strips were imaged using a laboratory scanner (Azure Biosystems 600), and band intensity was quantified using python.

### Monte Carlo Hypothesis Testing

Within the linear range of camera detection, particle intensities tend to convolve under aggregation (dimer intensities are the auto-convolution of the monomer distribution). Baseline particle distributions contain a mixture of monomer, dimer, trimer, and other higher order species. To model the baseline intensity distribution simply, we chose to approximate the distribution as a mixture of normal distributions (as normal distributions are closed under convolution). For naïve crosslinking, a particle will participate in j crosslink events according to a binomial distribution, and since *N* is large (> 1 million) and the probability tends to be small (< 0.1) we chose Poisson weightings for the monomer, dimer, etc. component distributions. Thus, we create a baseline distribution with the form: *B*(λ, μ, σ) = ∑_*i*=1_ Poisson(*i* − 1, λ)Normal(*i*μ, iσ). Here, λ captures the polydispersity of the baseline particles. Lower λ results in greater monomer. Next, we introduce a cross-linking rate *k*, ranging from 0.1 down to 1e-8. To “crosslink” the baseline distribution we convolve the baseline distribution with itself 4 times and create a new mixture distribution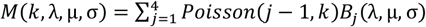, where *B*_*j*_(λ, μ, σ) is the j-th convolution of the baseline with itself. Only four self-convolutions were considered for since the components quickly drop off to bellow 1e-12.

To perform the hypothesis testing, 300 samples of size n (varying from 1e1 to 1e8) were drawn from the baseline *B*(λ, μ, σ) and crosslinked distribution *M*(*k*, λ, μ, σ). The log-likelihood ratio, Λ, was computed on both B and M samples. A α level of 0.05 is used on the baseline B samples, meaning a threshold on Λ is chosen such that only 5 percent of samples are larger. Next, the β of the threshold was computed to be the fraction of M samples with Λ below the threshold. If the power (1 − β) was below 0.95, then a larger sample size (n) is attempted until sufficient power is reached.

The crosslinking rate, *k*, is approximately the fraction of higher order species and is converted to concentration by assuming perfect crosslinking. For example, for k = 0.01, then 0.00995 of the initial particles will be crosslinked in higher order states, so assuming 100 million initial particles in 25 uL allows conversion to concentration. For imperfect crosslinking, for example a 1 pM binder, then using a simplified equilibrium analysis with excess capture ligand, would mean that each crosslinker has a > 99% probability of being bound, thus mildly shifting the scaling curves to the right.

### Statistical Analysis

Data analysis was performed using Python. Error bars represent standard error unless otherwise noted.

## Acknowledgements and Funding

All financial support for this work was provided by ilytica.

## Author Contributions

DBM and JTG wrote the manuscript. JTG proposed the general image-analytic framework. DBM conceived the machine learning approach and implemented both hybrid and end-to-end machine learning strategies. QH, AA, and HJW prepared nanoparticles for CRP and S.CoV-2 IgG, and also performed CRP experiments with conventional image analysis. HJW supervised CRP experiments and analysis performed by QH and AA. DBM performed the experiments, analysis, and ML training for S.CoV-2 IgG detection. DBM did the theoretical analysis of sensitivity scaling laws by particle count. QH, AA, and HJW performed work at Texas A&M University under contract with ilytica. All other work was performed at ilytica.

## Data and Material Availability

The datasets, code, and custom materials generated and/or analyzed in this study are not publicly available due to intellectual property considerations. They may be made available to qualified researchers or organizations upon reasonable request, subject to the execution of an appropriate legal agreement (e.g., non-disclosure agreement or material transfer agreement) between the requesting party and ilytica, LLC. Requests for access should be directed to

## Supplemental Information

**Supplementary Figure 2.**
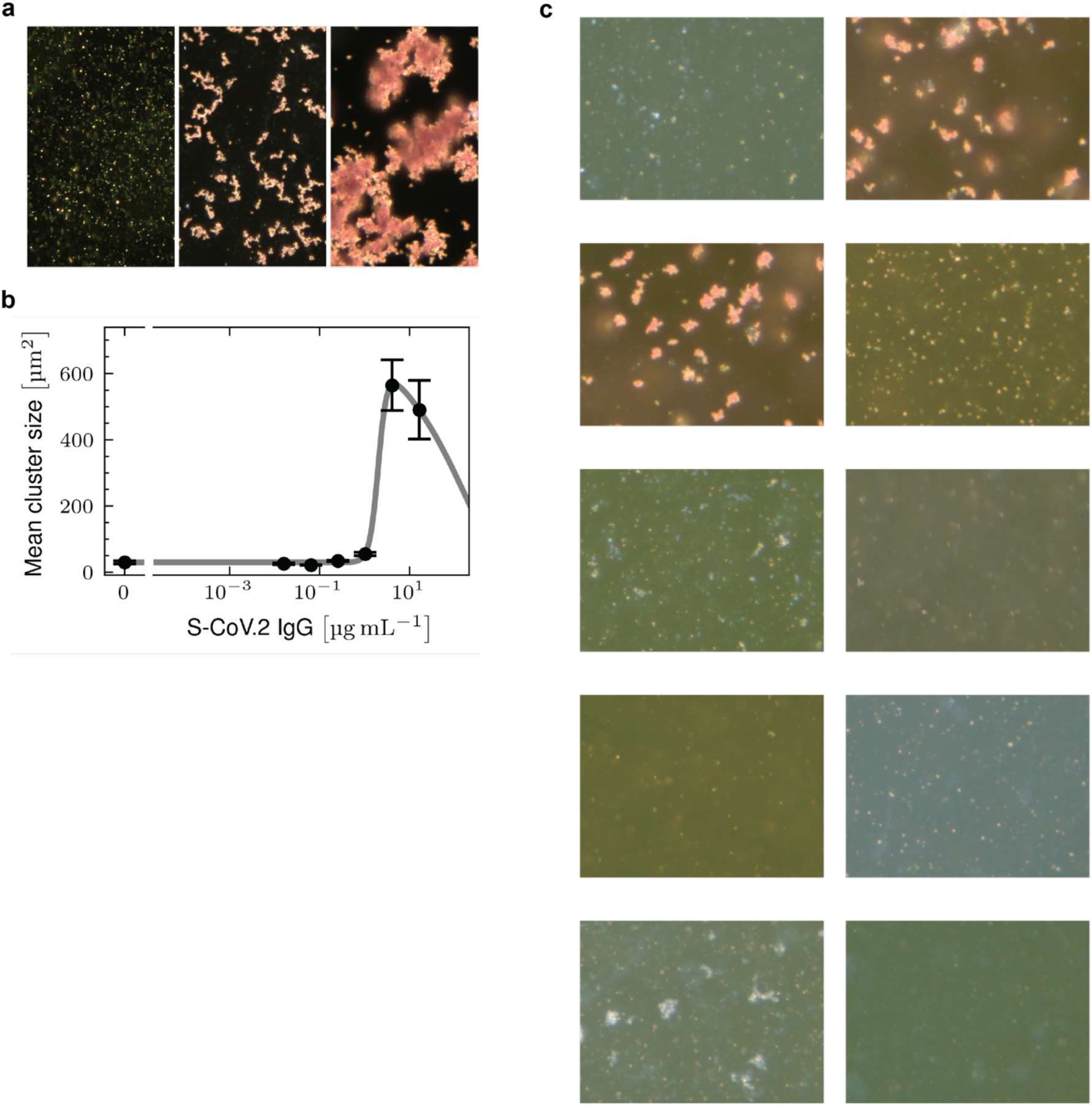
(a) Example images of particle aggregation in the presence of increasing antibody (Ab against S-CoV.2 RBD). 20x darkfield was used, (b) Quantification of cluster size as a function of S-CoV.2 IgG concentration, (c) Example images taken using 20x darkfield in open air microwells with notable optical artifacts.

## References

1. Rolando, J. C., Melkonian, A. V. & Walt, D. R. The Present and Future Landscapes of Molecular Diagnostics. Annu. Rev. Anal. Chem. 17, 459–474 (2024).

2. Rissin, D. M. et al. Single-molecule enzyme-linked immunosorbent assay detects serum proteins at subfemtomolar concentrations. Nat Biotechnol 28, 595–599 (2010).

3. Wu, A. H., Fukushima, N., Puskas, R., Todd, J. & Goix, P. Development and Preliminary Clinical Validation of a High Sensitivity Assay for Cardiac Troponin Using a Capillary Flow (Single Molecule) Fluorescence Detector. Clin. Chem. 52, 2157–2159 (2006).

4. Yelleswarapu, V. et al. Mobile platform for rapid sub–picogram-per-milliliter, multiplexed, digital droplet detection of proteins. Proc. Natl. Acad. Sci. 116, 4489–4495 (2019).

5. Yuan, L. et al. Digitizing Gold Nanoparticle-Based Colorimetric Assay by Imaging and Counting Single Nanoparticles. Anal. Chem. 88, 2321–2326 (2016).

6. Mao, C.-P. et al. Protein detection in blood with single-molecule imaging. Sci. Adv. 7, eabg6522 (2021).

7. Groves, J. T. Digital molecular assays. (2023).

8. USFDA. Recall Triage Cardiac Panel. https://www.accessdata.fda.gov/scripts/cdrh/cfdocs/cfRES/res.cfm?id=81652 (2009).

9. Shaw, J. L. V. Practical challenges related to point of care testing. Pr. Lab. Med. 4, 22–29 (2016).

10. Calarco, S. et al. Analytical performance of 17 commercially available point-of-care tests for CRP to support patient management at lower levels of the health system. PLOS ONE 18, e0267516 (2023).

11. Martiáñez-Vendrell, X., Skjefte, M., Sikka, R. & Gupta, H. Factors Affecting the Performance of HRP2-Based Malaria Rapid Diagnostic Tests. Trop. Med. Infect. Dis. 7, 265 (2022).

12. Laryea, E. T. & Nichols, J. H. Implementing Individualized quality control plans and managing risk at the point-of-care for molecular diagnostics. Expert Rev. Mol. Diagn. ahead-of-print, 1–7 (2023).

13. Chin, C. D., Linder, V. & Sia, S. K. Commercialization of microfluidic point-of-care diagnostic devices. Lab a Chip 12, 2118–2134 (2012).

14. Jackson, H. et al. MarkerDB 2.0: a comprehensive molecular biomarker database for 2025. Nucleic Acids Res. 53, D1415–D1426 (2024).

15. Plebani, M. The detection and prevention of errors in laboratory medicine. Ann. Clin. Biochem. 47, 101–110 (2009).

16. Plebani, M. Errors in clinical laboratories or errors in laboratory medicine? Clin. Chem. Lab. Med. (CCLM) 44, 750–759 (2006).

17. John, G. K., Favaloro, E. J., Austin, S., Islam, M. Z. & Santhakumar, A. B. From errors to excellence: the pre-analytical journey to improved quality in diagnostics. A scoping review. Clin. Chem. Lab. Med. (CCLM) 0, (2025).

18. Delianu, C. et al. Medical Staff Training - Quality Initiative to Reduce Errors in the Pre-Preanalytical Phase. Clin. Lab. 67, (2021).

19. Azzam, I., Pandori, M. & Sherych, L. Discontinue the Use of Antigen Testing in Skilled Nursing Facilities Until Further Notice. https://dpbh.nv.gov/uploadedFiles/dpbhnvgov/content/Resources/Directive%20to%20Discontinue%20Use%20of%20Antigen%20POC_10.02.2020_ADA_Compliant.pdf (2020).

20. Budd, J. et al. Lateral flow test engineering and lessons learned from COVID-19. Nat. Rev. Bioeng. 1, 13–31 (2023).

21. Enea, M., Leite, A., Franco, R. & Pereira, E. Gold Nanoprobes for Robust Colorimetric Detection of Nucleic Acid Sequences Related to Disease Diagnostics. Nanomaterials 14, 1833 (2024).

22. Kim, J. et al. Sensitive, Quantitative Naked-Eye Biodetection with Polyhedral Cu Nanoshells. Adv. Mater. 29, (2017).

23. Wu, H.-J. et al. Membrane-protein binding measured with solution-phase plasmonic nanocube sensors. Nat Methods 9, 1189 1191 (2012).

24. Baksh, M. M., Jaros, M. & Groves, J. T. Detection of molecular interactions at membrane surfaces through colloid phase transitions. Nature 427, 139–41 (2004).

25. Todd, J. et al. Ultrasensitive Flow-based Immunoassays Using Single-Molecule Counting. Clin. Chem. 53, 1990–1995 (2007).

26. Khalifian, S., Raimondi, G. & Brandacher, G. The Use of Luminex Assays to Measure Cytokines. J Invest Dermatol 135, 1–5 (2015).

27. Winter, E. M. & Groves, J. T. Surface Binding Affinity Measurements from Order Transitions of Lipid Membrane-Coated Colloidal Particles. Anal. Chem. 78, 174–180 (2006).

28. Liu, Y., Zhan, L., Qin, Z., Sackrison, J. & Bischof, J. C. Ultrasensitive and Highly Specific Lateral Flow Assays for Point-of-Care Diagnosis. ACS Nano 15, 3593–3611 (2021).

29. Burger, W. & Burge, M. J. Digital Image Processing, An Algorithmic Introduction. Texts Comput. Sci. (2022) doi:10.1007/978-3-031-05744-1.

30. He, K., Zhang, X., Ren, S. & Sun, J. Deep Residual Learning for Image Recognition. arXiv (2015) doi:10.48550/arxiv.1512.03385.

31. Koczula, K. M. & Gallotta, A. Lateral flow assays. Essays Biochem 60, 111–120 (2016).

32. Sato, K., Hosokawa, K. & Maeda, M. Rapid Aggregation of Gold Nanoparticles Induced by Non-Cross-Linking DNA Hybridization. J. Am. Chem. Soc. 125, 8102–8103 (2003).

33. Anker, J. N. et al. Biosensing with plasmonic nanosensors. Nat. Mater. 7, 442–453 (2008).

34. Sönnichsen, C., Reinhard, B. M., Liphardt, J. & Alivisatos, A. P. A molecular ruler based on plasmon coupling of single gold and silver nanoparticles. Nat Biotechnol 23, 741–745 (2005).

35. Li, T., Wu, X., Liu, F. & Li, N. Analytical methods based on the light-scattering of plasmonic nanoparticles at the single particle level with dark-field microscopy imaging. Analyst 142, 248–256 (2016).

36. Sriram, M. et al. A rapid readout for many single plasmonic nanoparticles using dark-field microscopy and digital color analysis. Biosens. Bioelectron. 117, 530–536 (2018).

37. Bennett, D. et al. Machine Learning Color Feature Analysis of a High Throughput Nanoparticle Conjugate Sensing Assay. Anal. Chem. 95, 6550–6558 (2023).

38. Berg, S. et al. ilastik: interactive machine learning for (bio)image analysis. Nat Methods 16, 1226–1232 (2019).

39. Falk, T. et al. U-Net: deep learning for cell counting, detection, and morphometry. Nat. Methods 16, 67–70 (2019).

40. Ronneberger, O., Fischer, P. & Brox, T. U-Net: Convolutional Networks for Biomedical Image Segmentation. Lect Notes Comput Sc 234–241 (2015) doi:10.1007/978-3-319-24574-4_28.

41. Cox, R. J. et al. Seroconversion in household members of COVID-19 outpatients. Lancet Infect. Dis. 21, 168 (2021).

42. Plumb, I. D. et al. Estimated COVID-19 vaccine effectiveness against seroconversion from SARS-CoV-2 Infection, March–October, 2021. Vaccine 41, 2596–2604 (2023).

43. Christenson, R. H. & Duh, S.-H. Methodological and Analytic Considerations for Blood Biomarkers. Prog. Cardiovasc. Dis. 55, 25–33 (2012).

44. Sugimoto, T. Monodispersed Particles. (Elsevier, 2019). doi:10.1016/c2012-0-02740-8.

